# Chromosome-level, haplotype-resolved genome assembly of the tanniferous forage legume big trefoil (*Lotus pedunculatus* Cav.) using CiFi

**DOI:** 10.64898/2026.09.09.749848

**Authors:** Alexander T. Pettersson, Yutang Chen, Tim Davalan, Pamela Nicholson, David Kopecky, Bruno Studer, Roland Kölliker

**Affiliations:** Molecular Plant Breeding, Institute of Agricultural Sciences, ETH Zurich, Universitätstrasse 2, 8092, Zurich, Switzerland; Next Generation Sequencing Platform, University of Bern, Bern, Switzerland; Institute of Experimental Botany of the Czech Academy of Sciences, Centre of Plant Structural and Functional Genomics, Olomouc, Czech Republic

## Abstract

Big trefoil (*Lotus pedunculatus* Cav.) is a perennial forage legume that thrives on acidic, low-fertility soils and produces condensed tannins that reduce enteric methanogenesis in ruminants. Despite this agronomic potential, genomic resources for the species remain scarce, and the existing haploid assembly does not resolve the two haplotypes of this outcrossing diploid species. Here we present a haplotype-resolved, chromosome-level reference genome for *L. pedunculatus* genotype Lusitano29 – the first plant genome assembled using CiFi, a long-read chromosome conformation capture method. We combined PacBio HiFi long reads with CiFi concatemers produced from *Dpn*II and *Hind*III libraries; *in silico* digestion and combinatorial pairing of the resulting monomers yielded 790.3 M and 10.3 M pseudo-paired contacts, respectively, enabling scaffolding and manual curation to chromosome level. The 991.1 Mb assembly resolves two phased haplotypes of 500 and 491 Mb, with 96.6% of the sequence anchored in twelve pseudo-chromosomes (six per haplotype). Telomeric repeats were detected at 19 of 24 pseudo-chromosome ends, and no structural errors were detected (scaffold N50 73.8 Mb; consensus QV 64.7; *k*-mer completeness 99.4%; genome-mode BUSCO completeness 97.0%; CRAQ S-AQI 100.0). Annotation supported by PacBio Iso-Seq full-length transcripts predicted 38,069 and 36,484 protein-coding genes in haplotypes 1 and 2, respectively (protein-mode BUSCO completeness 96.5%), indicating a high completeness of annotated genes. This genome assembly provides a foundation for allele-aware trait dissection of proanthocyanidin biosynthesis, comparative genomics in *Lotus*, and population genomics and genomics-assisted breeding in *L. pedunculatus*.

## Background & Summary

The genus *Lotus* encompasses a diverse group of species widely utilized as forage crops and biological models^1,2^. Within *L*. sect. *Lotus*, big trefoil (*Lotus pedunculatus* Cav.) is a perennial forage legume distinguished from close relatives such as the model legume *Lotus japonicus* K. Larsen by, among other traits, its stoloniferous growth habit^3^. *L. pedunculatus* is of increasing agronomic interest not only because it remains productive in acidic and low-fertility soils where other forage crops fail^4^, but also because condensed tannins in its biomass can favorably shape the rumen microbiome of grazing livestock, reducing methanogenesis and increasing protein uptake^5^. *L. pedunculatus* is naturally diploid with a basic chromosome number of six (2n = 2x = 12), but artificial autotetraploids have been induced by chromosome doubling, including the hybrid cultivar ‘Grasslands Maku’, which was bred for improved biomass yield and vigor^6^.

Despite this great potential as a forage crop, genomic resources for *L. pedunculatus* remain limited. The Darwin Tree of Life Project has generated a haploid chromosome-level assembly (`GCA_965212415.1`); however, as a collapsed haploid representation, it does not resolve the two haplotypes and thus captures only part of the sequence diversity of the diploid genome. Therefore, a haplotype-resolved chromosome-level assembly is needed for this heterozygous diploid species to reduce reference bias inherent to collapsed haploid representations. Such a resource will enable allele-aware population genomics, together with comparative and breeding applications in *L. pedunculatus* and the genus *Lotus* more broadly, particularly given that large structural variants are known to segregate within *Lotus* species^7^.

Haplotype-resolved assemblies are typically phased and scaffolded with chromosome conformation capture data, for which methods such as Hi-C are widely used. However, short-read-based approaches frequently fail to map uniquely across repetitive and low-complexity regions, such as centromeres and segmental duplications^8^. CiFi addresses these limitations by coupling chromosome conformation capture with PacBio HiFi long-read sequencing, yielding sequencing reads of several kilobases composed of multiple ligated, spatially proximate monomers. This not only improves read mapping across repetitive regions, but also improves haplotype phasing and reduces input requirements by orders of magnitude relative to conventional Hi-C and other long-read chromatin conformation capture methods, enabling chromosome-level, haplotype-resolved assemblies from as few as tens of thousands of cells^8^.

Here, we present the first haplotype-resolved, chromosome-level genome assembly for *L. pedunculatus* genotype Lusitano29, generated using PacBio HiFi sequencing and CiFi chromosome conformation capture^8^. While CiFi has previously been demonstrated on human cell lines and single insects^8^, this work aims to extend the method to a plant genome. We anticipate that this high-quality genome assembly will support comparative and evolutionary studies within the genus *Lotus*, including synteny-based reconstruction of chromosome evolution against the model *L. japonicus*. We also expect it will enable trait dissection of condensed-tannin biosynthesis, from annotation of the proanthocyanidin pathway to investigation of allele-specific expression in proanthocyanidin genes. Finally, we anticipate this resource will assist breeding efforts in this outcrossing forage, including genomic selection for yield and persistence and marker-assisted introgression of traits^9^.

## Methods

### Plant material

Seeds of *Lotus pedunculatus* genotype Lusitano29 (IPK accession LOT 29; IPK Gatersleben, Germany; DOI: `10.25642/I PK/GBIS/64418`), originating from Portugal, were germinated and plants were cultivated at Aarhus University (Aarhus, Denmark). A single mature individual was transferred to ETH Zurich, where clonal cuttings were established and maintained under greenhouse conditions (16 h light / 8 h dark, 20 °C).

### Genome size estimation and karyotyping

Genome size was estimated by flow cytometry following Galbraith et al.^10^. Nuclear suspensions were prepared by simultaneously chopping 50 mg of leaf tissue from *L. pedunculatus* and the internal reference standard *Solanum lycopersicum* L. ‘Stupicke polni tyckove rane’ (2C = 1.96 pg)^11^ in 0.5 mL Otto I solution^12^ (0.1 M citric acid, 0.5% v/v Tween 20). The suspension was filtered through a 42-*µ*m nylon mesh and stained with 1 mL Otto II solution (0.4 M Na_2_HPO_4_*·* 12H_2_O) supplemented with 2 mg/mL *β*-mercaptoethanol, 50 *µ*L/mL propidium iodide and 50 *µ*L/mL RNase IIA. Samples were stained for 10 min at room temperature and analysed using a CyFlow flow cytometer (Partec, Münster, Germany) equipped with a 532-nm laser. At least 5,000 nuclei were analysed per sample and only histograms with a G0/G1 peak coefficient of variation below 3.0% were accepted. The analysis was repeated on three separate days.

For karyotyping, young roots were collected from hydroponically cultivated cuttings of Lusitano29 and pretreated in ice-cold distilled water for 24 h. Root tips were excised and immediately fixed in Farmer’s fixative (3:1 v/v; absolute ethanol *>*99.8% : glacial acetic acid *>*99.8%) and incubated at 37 °C for seven days. Fixed root tips were stained in 1% acetocarmine for 2 h and squashed in a drop of 45% acetic acid on clean microscope slides. Slides were frozen on a dry-ice block for 1 h, then counterstained with 1.5 *µ*g/mL 4^*′*^,6-diamidino-2-phenylindole in Vectashield antifade solution (Vector Laboratories, Newark, CA, USA). Slides were examined under an Axio Imager Z.2 Zeiss microscope (Carl Zeiss Microscopy, Jena, Germany) equipped with a Cool Cube 1 camera (Metasystems, Altlußheim, Germany). Images were captured and processed using ISIS v.5.4.7 (Metasystems) and Adobe Photoshop (Adobe, San Jose, CA, USA).

### HiFi whole-genome sequencing

Young leaves were harvested from clonal cuttings of Lusitano29, flash-frozen in liquid nitrogen, ground to a fine powder under liquid nitrogen and stored at −80 °C until shipment on dry ice to the Next Generation Sequencing Platform (NGSP), University of Bern, Switzerland.

Nuclei were isolated from the powdered leaf tissue according to the PacBio procedure *Isolating nuclei from plant tissue using LN2 disruption* (PacBio, 102-574-800, Rev. 04, January 2025). High-molecular-weight (HMW) DNA was subsequently extracted from the isolated nuclei using a cetyltrimethylammonium bromide (CTAB)-based method^13–16^. DNA concentration, fragment size distribution and purity were assessed using a Qubit 4 Fluorometer with the Qubit dsDNA HS and BR Assay Kits (Thermo Fisher Scientific, Q32851 and Q32850), an FEMTO Pulse system with the Genomic DNA 165 kb Kit (Agilent, FP-1002-0275) and a DeNovix DS-11 UV-Vis spectrophotometer, respectively.

HMW DNA was used to prepare PacBio SMRTbell libraries according to *Preparing whole genome and metagenome libraries using SMRTbell prep kit 3*.*0* (PacBio, 102-166-600, Rev. 06, January 2025), with the modifications described below. HMW DNA was not treated with a short-read eliminator and was instead directly sheared using a Megaruptor 3 system (Diagenode, B06010003) with a Megaruptor 3 Shearing Kit (Diagenode, E07010003). Sheared DNA was purified using 1× SMRTbell cleanup beads and fragment size distributions were assessed using the FEMTO Pulse system as described above. Samples with fragment sizes predominantly within the 13–20 kb range were carried forward for library preparation.

Subsequent library preparation followed the manufacturer’s protocol and included end repair and A-tailing, ligation of barcoded overhang adapters, AMPure PB bead purification and nuclease treatment. As a modification to the standard workflow, gel-based size selection was performed to remove fragments *<*10 kb according to the PacBio technical note *Gel Cassette Size Selection Methods for WGS HiFi Libraries*, using the protocol for the Sage Science BluePippin system (PacBio, 102-326-503, Rev. 02, May 2024). Final SMRTbell libraries were assessed for concentration and fragment size distribution using Qubit and FEMTO Pulse, respectively, as described above.

Libraries were prepared for sequencing according to the SMRT Link Sample Setup workflow (SMRT Link v25). Using components from the Revio SPRQ Polymerase Kit (PacBio, 103-496-900) and cleanup beads (PacBio, 102-158-300), the standard PacBio sequencing primer was annealed to the SMRTbell libraries, followed by binding of Revio DNA polymerase and bead-based purification of the resulting polymerase-bound complexes. Revio sequencing control DNA was diluted and added to the complexes before loading onto a thawed Revio SPRQ sequencing plate (PacBio, 102-326-552).

The Revio instrument deck was prepared according to the SMRT Link workflow, including sequencing plates, pipette tips and Revio SMRT Cell trays containing four SMRT Cell 25M sequencing cells (PacBio, 102-202-200). Libraries were loaded at an on-plate concentration of 300 pM using adaptive loading. SMRT sequencing was performed on a PacBio Revio system with a 30-h movie time. Basic data processing, including demultiplexing of barcoded SMRTbell libraries, was performed using SMRT Link v25.

All procedures following shipment of the liquid-nitrogen-ground leaf material, including nuclei isolation, HMW DNA extraction, library preparation, quality control and sequencing, were performed at NGSP.

### CiFi chromosome conformation capture

Nuclei were isolated from 1 g of the same liquid-nitrogen-ground Lusitano29 leaf material used for HiFi whole-genome sequencing, following the nuclei isolation procedure described above. CiFi chromosome conformation capture was performed using a protocol adapted from the method described by McGinty et al.^8^.

Plant nuclei were resuspended in 1 mL of 1× phosphate-buffered saline (PBS) and crosslinked with 2% formaldehyde for 10 min at room temperature. Crosslinking was quenched with 92 *µ*L of Stop Solution 1 (Arima Genomics, A311049) for 15 min on ice. Nuclei were collected by centrifugation at 3,000*× g* for 5 min at room temperature, washed in 1 ×PBS and stored as pellets at −80 °C until further processing.

Nuclei were thawed on ice and resuspended in 50 *µ*L of 10× Protease Inhibitor Cocktail (Promega, G6521) and 500 *µ*L of cold permeabilization buffer (10 mM Tris-HCl, pH 8.0, 10× mM NaCl, 0.2% IGEPAL CA-630). Following incubation on ice for 15 min, the sample was split 50:50 into two separate tubes – one designated for *Dpn*II digestion and the other for *Hind*III digestion – and nuclei were collected by centrifugation at 500*× g* for 10 min.

Pellets were washed and resuspended in the corresponding restriction-enzyme buffer: *Dpn*II buffer (NEB, B0543S) for *Dpn*II or NEBuffer r2.1 (NEB, B6002S) for *Hind*III. SDS was added to a final concentration of 0.1%, followed by incubation at 65 °C for 10 min with mixing at 300 rpm. Samples were cooled on ice, Triton X-100 was added to a final concentration of 1% and samples were incubated for a further 10 min on ice. A 30 *µ*L aliquot of each sample was retained and frozen as a non-digested control.

Chromatin was digested with *Dpn*II (NEB, R0543L) or *Hind*III (NEB, R0104L) in the respective parallel reactions, each at a final enzyme concentration of 1 U/*µ*L in 1× restriction buffer, for 18 h at 37 °C with mixing at 300 rpm. Restriction enzymes were subsequently heat-inactivated for 20 min at 65 °C for *Dpn*II and 80 °C for *Hind*III and samples were immediately cooled on ice. A 50-*µ*L aliquot from each digestion was retained and frozen as a non-ligated control.

Proximity ligation was performed in a final volume of 1 mL containing 1 ×T4 DNA Ligase Buffer (NEB, B0202S), 1 mg bovine serum albumin (BSA; Thermo Fisher Scientific, AM2616) and 20,000 U T4 DNA Ligase (NEB, M0202L). Samples were incubated for 6 h at 16 °C with mixing at 300 rpm.

Crosslinks were subsequently reversed by addition of Proteinase K (100 *µ*L of a 20 mg/mL solution; 2 mg total), SDS (100 *µ*L of a 10% solution; 0.5% final) and Tween-20 (500 *µ*L of a 20% solution; 5% final), followed by incubation for 18 h at 56° C.

DNA from the CiFi samples and the non-digested and non-ligated controls was purified by phenol–chloroform extraction. Briefly, overnight samples were split into 2× 1 mL aliquots and control samples were brought to 200 *µ*L; all samples were then extracted sequentially with equal volumes of phenol:chloroform:isoamyl alcohol (25:24:1) and chloroform:isoamyl alcohol (24:1), with phase separation by centrifugation at 16,000 *×g* for 5 min. DNA was precipitated with 0.75 M ammonium acetate, glycogen and 2.5 volumes of 100% ethanol, followed by centrifugation at 16,000*× g* for 20 min at 4 °C.

Pellets were washed twice with 80% ethanol, briefly air-dried and resuspended in PacBio Elution Buffer (PacBio, 101-633-500; 10 *µ*L for controls, 50 *µ*L for experimental samples) at room temperature overnight. DNA concentration and fragment size distribution were assessed using a Qubit 4 Fluorometer with the Qubit dsDNA HS Assay Kit (Thermo Fisher Scientific, Q32851) and an FEMTO Pulse system with the Genomic DNA 165 kb Kit (Agilent, FP-1002-0275), respectively.

Low-input amplification and SMRTbell library preparation were performed on this CiFi-extracted DNA according to the PacBio procedure *Amplifying genomic DNA for SMRTbell library preparation and HiFi sequencing* (PacBio, 103-648-000, Rev. 04, July 2025). Briefly, 1–50 ng of CiFi DNA was processed according to Option 1 of the protocol, using DNA shearing with the Megaruptor 3 system (Diagenode). Sheared DNA was purified using 1 ×SMRTbell cleanup beads (PacBio, 102-158-300), followed by end repair, A-tailing and ligation of amplification adapters. Adapter-ligated DNA was PCR-amplified and purified using a 1× bead cleanup.

Amplified and barcoded DNA was subsequently used as input for SMRTbell library preparation. Libraries were size-selected using AMPure PB beads (PacBio, 100-265-900) and library concentration and fragment size distribution were assessed using Qubit and FEMTO Pulse, respectively.

Libraries were prepared for sequencing according to the SMRT Link Sample Setup workflow (SMRT Link v25). Using components from the Revio SPRQ Polymerase Kit (PacBio, 103-496-900) and cleanup beads (PacBio, 102-158-300), the standard PacBio sequencing primer was annealed to the SMRTbell libraries, followed by binding of Revio DNA polymerase and bead-based purification of the polymerase-bound complexes. Revio sequencing control DNA was added before loading onto a thawed Revio SPRQ sequencing plate (PacBio, 102-326-552).

The Revio instrument deck was prepared according to the SMRT Link workflow using Revio SMRT Cell trays containing four SMRT Cell 25M sequencing cells (PacBio, 102-202-200). Libraries were loaded at an on-plate concentration of 300 pM using adaptive loading. SMRT sequencing was performed on a PacBio Revio system with a 30-h movie time. Basic data processing was performed using SMRT Link v25, including demultiplexing of indexes introduced using the Twist Universal Adapter System (Twist Bioscience, 101307–101311) and SMRTbell barcodes, followed by marking of PCR duplicates.

All procedures following receipt of the liquid-nitrogen-ground leaf material, including nuclei isolation, CiFi chromosome conformation capture, DNA purification, library preparation, quality control, sequencing and basic data processing, were performed at NGSP.

### Iso-Seq transcriptome sequencing

Fourteen samples representing seven tissues (root, stem, flower, leaf, whole inflorescence, petal and reproductive whorl) were collected from Lusitano29, flash-frozen in liquid nitrogen and stored at− 80 °C. Frozen tissues were ground to a fine powder under liquid nitrogen and total RNA was extracted using the NucleoSpin RNA Plus Kit (MACHEREY-NAGEL, Düren, Germany) according to the manufacturer’s protocol, with *β*-mercaptoethanol added to the lysis buffer to a final concentration of 1% (v/v). RNA was eluted in 30 *µ*L and treated with DNase I using the DNA-free DNA Removal Kit (Invitrogen, Waltham, MA, USA) according to the manufacturer’s instructions. RNA integrity and concentration were initially assessed using the RNA ScreenTape assay on a TapeStation system (Agilent Technologies). Four samples with an RNA integrity number *>* 7.5 (flower, root, leaf and whole inflorescence) were shipped on dry ice to NGSP.

At NGSP, RNA concentration and integrity were reassessed using a Qubit 4 Fluorometer with the Qubit RNA HS Assay Kit (Thermo Fisher Scientific) and an FEMTO Pulse system with the Ultra Sensitivity RNA Kit (Agilent Technologies), respectively. Samples with an RNA quality number *>* 7 were pooled in equal RNA mass and used as input for PacBio Kinnex full-length RNA library preparation according to the PacBio procedure *Preparing Kinnex libraries using the Kinnex full-length RNA kit* (PacBio, 103-238-700, Rev. 06, December 2024). Briefly, 300 ng of total RNA per sample was reverse-transcribed to generate first-strand cDNA, followed by PCR amplification and barcoding using barcoded Iso-Seq primers. Equal masses of barcoded cDNA were pooled to a total input of 55 ng for Kinnex amplification. Eight parallel Kinnex PCR reactions were performed to introduce direction-specific Kinnex segmentation sequences. The amplified cDNA products were subsequently processed with Kinnex enzyme, ligase and barcoded Kinnex terminal adapters to concatenate cDNA segments into linear arrays. Following nuclease treatment and bead-based purification, the final Kinnex library was quantified using a Qubit 4 Fluorometer with the Qubit dsDNA HS Assay Kit (Thermo Fisher Scientific, Q32854) and fragment size distribution was assessed using an FEMTO Pulse system with the Genomic DNA 165 kb Kit (Agilent, FP-1002-0275).

The Kinnex SMRTbell library was prepared for sequencing by annealing the Kinnex sequencing primer, binding Revio DNA polymerase and performing bead-based cleanup using components from the Revio SPRQ Polymerase Kit (PacBio, 103-496-900) and SMRTbell cleanup beads (PacBio, 102-158-300). The polymerase-bound library was prepared for loading according to the SMRT Link v25 Sample Setup workflow, including calculation of the final loading dilution and addition of Revio sequencing control DNA.

The prepared library was loaded onto a Revio SPRQ sequencing plate (PacBio, 103-504-900). The Revio instrument deck was configured according to SMRT Link v25 using the required sequencing plate, pipette tips and a Revio SMRT Cell tray containing four SMRT Cell 25M sequencing cells (PacBio, 102-202-200). The library was loaded at an on-plate concentration of 160 pM using adaptive loading and SMRT sequencing was performed on a PacBio Revio system with a 30-h movie time.

Primary and subsequent data processing were performed using SMRT Link v25. Kinnex reads were demultiplexed and segmented on-instrument into S-reads. The Iso-Seq pipeline in SMRT Link v25 was then used for cDNA demultiplexing, poly(A) tail removal and identification of full-length transcript reads.

All procedures following RNA extraction, including RNA quality control, Kinnex full-length RNA library preparation, SMRTbell library preparation, sequencing and basic data processing, were performed at NGSP.

### Pre-assembly analyses

Prior to assembly, the sequencing reads were converted to analysis-ready formats, assessed for quality and contamination and used to characterize genome properties such as ploidy, genome size and heterozygosity. HiFi and CiFi `.bam` files were converted to compressed FASTQ format using SAMtools v1.23^17^ (`samtools fastq`). To verify sequencing output and quality, read length, base quality and GC content summaries for the HiFi whole-genome sequencing and CiFi datasets were generated using Seqkit v2.12.0^18^ (`seqkit stats`).

To detect potential contamination in the sequencing data, HiFi whole-genome sequencing reads were screened with Kraken2 v2.17.1^19^ (`kraken classify`) against the broad-taxon PlusPFP *k*-mer database (v2026-02-26).

To estimate ploidy, heterozygosity and haploid genome size prior to assembly, *k*-mer databases were built from the HiFi reads with FastK v1.2^20^ using a *k*-mer size of 41 and analysed with Smudgeplot v0.5.3^21^ and GenomeScope2 v2.1.0^21^.

To generate pseudo-paired contact data for assembly and scaffolding, CiFi concatemers were processed using a custom Python script. Briefly, Biopython v1.87^22^ was used to digest concatemers into all possible monomers; single-monomer concatemers were discarded and monomers shorter than 50 bp were excluded. For each concatemer, all possible pairwise monomer combinations were created using the itertools module of the Python standard library v3.13.4 and split into separate pseudo-paired read files.

### Genome assembly and scaffolding

To reduce computational demand while retaining sufficient coverage for haplotype-resolved assembly, the HiFi whole-genome sequencing data were randomly downsampled at the read level to 40× coverage of the diploid genome using Seqkit v2.12.0^18^ (`seqkit sample`) prior to assembly of the nuclear genome. The genome was assembled with hifiasm v0.25.0-r726^23–25^ using the downsampled HiFi reads and the CiFi pseudo-paired contact data as input (`-h1 dpnII_R1.fq.gz,hindIII_R1.fq.gz`;`-h2 dpnII_R2.fq.gz,hindIII_R2.fq.gz`). Default parameters were used, except that the genome size estimate was set to 541 Mb based on the flow cytometry estimate (`-hg-size 541m`) and the canonical plant telomeric repeat motif was provided for telomere identification during assembly (`-telo-m CCCTAAA`).

Contig sequences from the phased haplotype 1 (H1) and haplotype 2 (H2) assembly graphs were converted to FASTA format and concatenated into a diploid reference (H1+H2), ensuring competitive read mapping and simultaneous scaffolding of both haplotypes. CiFi pseudo-paired reads were aligned to the diploid assembly using BWA v0.7.19^26^ (`bwa mem -5SP`). Alignments were filtered with SAMtools v1.23^17^ (`samtools view -F 2316`), thus retaining only properly paired primary alignments. *Dpn*II and *Hind*III alignments were then merged. The resulting alignment file served as the input for both scaffolding approaches described below.

To obtain an automated chromosome-scale scaffolding for comparison with manual curation, the assembly was first scaffolded with HapHiC v1.0.7^27^ in pipeline mode (`haphic pipeline`), requesting twelve clusters corresponding to the expected chromosome number (`-nchrs 12`) and providing both haplotype assembly graphs (`-gfa`). The restriction motifs of the two enzymes used for CiFi library preparation were supplied for contact filtering (-RE GATC,AAGCTT); the maximum Markov clustering inflation value was set to 3 (`-max_inflation 3`) and the minimum group length to 35 (`-min_group_len 35`).

For manual scaffolding and error correction, a contact map was generated with PretextMap v0.2.4^28^ from the same pseudo-paired read alignment file used for HapHiC. Scaffolding was then performed interactively in PretextView v1.0.7^29^, where contigs were visually inspected, ordered, oriented, and joined into chromosome-scale scaffolds, while chimeric misjoins were split and unplaced sequences were integrated based on patterns of chromatin interactions. Following manual curation, the pseudo-paired reads were re-mapped to the curated assembly and the final contact map was visualized using the `HapHiC plot` module of HapHiC v1.0.7^27^. Curated scaffolds and unplaced contigs were separated into their respective H1 and H2 haplotype assemblies. Scaffolds and unplaced sequences in each haplotype assembly were then ordered by size and scaffolds renamed `chr_1` to `chr_6`. Assembly quality was evaluated across multiple tools using the PAQman v1.2.0^30^ toolkit. Contiguity metrics and length statistics were computed using QUAST v5.3.0^31^. Gene-space completeness was assessed with BUSCO v6.0.0^32,33^ in genome mode against the `fabales_odb12.2025-07-01` lineage dataset. Reference-free *k*-mer completeness and consensus quality values were calculated using Merqury v1.3^34^ and meryl v1.3^34^ using the downsampled HiFi read set. Finally, regional and structural assembly errors were profiled at single-base resolution using CRAQ v1.0.9^35^ supplemented with the downsampled HiFi reads in combination with bedtools v2.31.1^36^. Genome alignments required for error evaluation were performed with minimap2 v2.30^37^ and BWA v0.7.19^26^.

### Gene annotation

Structural gene annotations for both H1 and H2 assemblies were established through a hybrid strategy combining extrinsic transcript and homology evidence with deep-learning *ab initio* prediction. Evidence-based annotation was first performed with EviAnn v2.0.5^38^ (maximum intron size 50 kb), utilizing full-length Iso-Seq transcripts, *Lotus japonicus* coding sequences (v1.3) from Lotus Base^39^, and 305,615 Hologalegina RefSeq proteins (NCBI, downloaded 2026-05-24). To capture genes that were not expressed in the sampled tissues or genes that lacked sufficient protein homology, genome-wide *ab initio* predictions were generated with the Helixer^40^ web service (accessed 2026-05-27, https://www.plabipd.de/helixer_main.html) using the land-plant lineage model. The evidence-based EviAnn annotations were complemented with Helixer predictions using the `agat_sp_complement_annotations.pl` script from AGAT v0.8.0^41^, such that Helixer models were incorporated only where they did not overlap an existing gene annotated by EviAnn, thus leaving all EviAnn gene models unchanged. Functional annotation of predicted proteins across both haplotypes was carried out using EggNOG-mapper v2.1.13^42^ and InterProScan v5.77 (data release 108.0)^43,44^. Using EggNOG-mapper, sequences were queried using MMseqs2 (`-m mmseqs`, `-itype proteins`), with orthology assignment restricted to Viridiplantae (`-tax_scope 33090`, `-target_taxa 33090`) and all GO evidence codes retained (`-go_evidence all`). In InterProScan, protein sequences (`-seqtype p`) were analysed with precalculated lookup disabled (`-disable-precalc`) and both InterPro entry lookup (`-iprlookup`) and GO term mapping (`-goterms`) enabled.

### Transposable element annotation

Transposable elements and repetitive DNA sequences were identified and annotated across both haplotypes independently using EDTA v2.3.0^45–49^. *De novo* repeat libraries were generated without an external curated reference by specifying generic plant parameters (`-species others`). The complete EDTA pipeline was executed (`-step all`), enabling sensitive repeat detection with RepeatModeler to recover non-canonical or divergent elements (`-sensitive 1`), followed by whole-genome annotation (`-anno 1`) and consistency evaluation (`-evaluate 1`).

### Visualization of genomic features

To visualize genome-wide features and structural characteristics of the assembly, circos plots and summary tracks were generated using PyCirclize v1.10.1. Telomeric regions were identified by scanning 1 Mb pseudo-chromosome termini for the canonical plant telomere repeat motif (`CCCTAAA` at 5^*′*^ ends, `TTTAGGG` at 3^*′*^ ends) using approximate matching (Hamming distance*≤* 1). GC content was calculated in 20 kb sliding windows with a 5 kb step size and plotted as deviation from the global mean. Gene density and the proportion of transposable elements were calculated in 1 Mb windows with a 500 kb step size using feature midpoints for genes and base-pair coverage for transposable elements, with overlapping transposable element intervals merged to avoid double-counting. Sequence-level synteny between the two haplotypes was evaluated by aligning the H2 pseudo-chromosomes against H1 using FastGA v1.5^50^ with a sequence identity threshold of 80% and a minimum alignment length of 2 kb. The resulting alignments were inspected to evaluate inter-chromosomal structural integrity, and subsequently filtered to only retain alignments between pairs of homologous pseudo-chromosomes.

## Data Records

All sequencing data, genome assemblies, and annotations generated in this study have been deposited in the European Nucleotide Archive (ENA) at EMBL-EBI under BioProject accession number PRJEB120729^51^.

The PacBio HiFi whole-genome sequencing and CiFi chromosome conformation capture reads (*Dpn*II and *Hind*III libraries) from leaf tissue of *Lotus pedunculatus* genotype Lusitano29 have been deposited under BioSample accession ERS30735524. The PacBio Kinnex Iso-Seq full-length transcriptome sequencing reads (segmented full-length S-reads) derived from pooled floral, foliar, and root tissues have been deposited under BioSample accession ERS30735525 (Table 1).

**Table 1.**
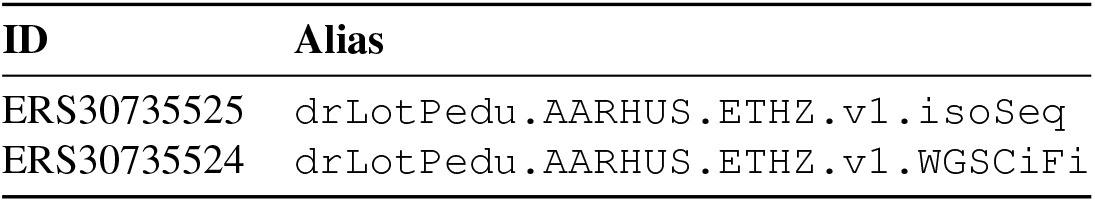
BioSamples registered at the European Nucleotide Archive for this study under project PRJEB120729. Both samples derive from the single sequenced individual of *L. pedunculatus* genotype Lusitano29 (NCBI taxon 347994). Sample ERS30735524 comprises the HiFi whole-genome sequencing (two runs) and CiFi (*Dpn*II and *Hind*III) libraries prepared from ground leaf tissue. Sample ERS30735525 comprises the Iso-Seq full-length transcriptome library prepared from an equimolar pool of flower, root, leaf, and whole-inflorescence RNA. The deposited Iso-Seq reads are the on-instrument segmented full-length S-reads.

The chromosome-level phased genome assemblies of *L. pedunculatus* genotype Lusitano29 for H1 (LPED_H1)^52^ and H2 (LPED_H2)^53^ have been deposited in ENA under analysis accession numbers ERZ29887158 and ERZ29887163, respectively. The structural gene annotations, transposable element annotations, and functional annotations (EggNOG-mapper and InterProScan) for both H1 and H2 have been deposited in ENA under analysis accessions ERZ29887156–ERZ29887165 (Table 2).

**Table 2.** Analysis objects submitted to the European Nucleotide Archive for *Lotus pedunculatus* genotype Lusitano29. All analyses were submitted under study ERP200841 (project PRJEB120729) and are linked to BioSample ERS30735524 (Table 1); structural gene annotation additionally used Iso-Seq data from BioSample ERS30735525.

| ID | Alias |
| --- | --- |
| ERZ29887156 | LPED_H1_annotation_gff_v3 |
| ERZ29887157 | LPED_H1_edta_gff_v3 |
| ERZ29887159 | LPED_H1_emapper_tab_v3 |
| ERZ29887160 | LPED_H1_interproscan_tab_v3 |
| ERZ29887158 | LPED_H1_assembly_fasta_v3 |
| ERZ29887161 | LPED_H2_annotation_gff_v3 |
| ERZ29887162 | LPED_H2_edta_gff_v3 |
| ERZ29887164 | LPED_H2_emapper_tab_v3 |
| ERZ29887165 | LPED_H2_interproscan_tab_v3 |
| ERZ29887163 | LPED_H2_assembly_fasta_v3 |

## Technical Validation

### Sequencing output

Sequencing yield and quality metrics for all libraries are summarized in Table 3. The HiFi whole-genome sequencing library was sequenced in two runs: a dedicated Revio SMRT cell yielding 137.4 Gb and a shared SMRT cell loaded at half-cell output equivalent yielding 78.4 Gb, for a combined 215.8 Gb (14.2 M reads; mean read length ~ 15.1 kb), corresponding to 199.4-fold coverage of the 1,082 Mb diploid genome. Median base qualities of Q36–Q37 with *≥* 95% of bases at Q30 or above meet expectations of highly accurate PacBio HiFi reads. CiFi sequencing yielded 53.0 Gb (8.9 M reads; mean read length 6.0 kb) for the *Dpn*II library and 15.5 Gb (2.4 M reads; mean read length 6.5 kb) for the *Hind*III library, with median base qualities of Q46, reflecting the higher per-base accuracy achieved by use of the AmpliFi protocol. Iso-Seq sequencing yielded 81.5 Gb of polymerase reads, from which on-instrument read segmentation produced 41.6 M full-length S-reads for use as transcript evidence in gene annotation. All data were delivered via the PacBio SMRT Link software.

**Table 3.** Sequencing yield and quality metrics across all libraries for *Lotus pedunculatus* genotype Lusitano29. Yield, read count, mean read length, and base quality were derived from PacBio primary analysis reports. Iso-Seq metrics refer to polymerase reads prior to on-instrument segmentation.

| Library | Yield [Gb] | Reads [M] | Mean length [kb] | Bases ≥ Q30 [%] |
| --- | --- | --- | --- | --- |
| HiFi WGS, run 1 | 137.4 | 9.1 | 15.1 | 95.1 |
| HiFi WGS, run 2 | 78.4 | 5.2 | 15.2 | 95.7<sup>a</sup> |
| CiFi *DpnII* | 53.0 | 8.9 | 6.0 | 97.5<sup>a</sup> |
| CiFi *HindIII* | 15.5 | 2.4 | 6.5 | 97.5<sup>a</sup> |
| Iso-Seq | 81.5 | 5.3 | 15.1 | 96.1<sup>a</sup> |

<sup>a</sup> Median base quality calculated from the whole flowcell.

### Contamination assessment

Kraken2 screening of the HiFi reads indicated negligible contamination. Of the 14.2 M HiFi whole-genome sequencing reads screened with Kraken2^19^, 99.91% were classified as eukaryotic and 98.32% were assigned to the genus *Lotus*. The remaining signal was dominated by reads assigned to other genera of the inverted repeat-lacking clade of legumes (1.12%; 159,977 reads), consistent with limited taxonomic resolution of the *k*-mer database among closely related legumes rather than true contamination. Trace assignments to human (0.18%; 26,114 reads) are attributable to routine laboratory handling and assignments to fungi, bacteria and viruses were negligible (*≤* 0.04%). Therefore, no evidence of significant biological contamination was identified in the sequencing data.

### Genome size and ploidy estimation

Karyotyping of root-tip preparations from clonal cuttings of Lusitano29 confirmed a chromosome number of 2*n* = 2*x* = 12, with two notably large chromosomes (Fig. 1a); these may correspond to the two largest assembled scaffolds (128.4 and 133.5 Mb; Fig. 4). Flow cytometry with *Solanum lycopersicum* as internal standard estimated the genome size at 541 Mb (Fig. 1b). GenomeScope2^21^ analysis of 41-mers under diploid assumptions estimated a haploid genome size of 487.8 Mb with a heterozygosity of 1.08% and a unique-sequence fraction of 66.7% (Fig. 1c), with clearly separated heterozygous (~ 42 ×) and homozygous (~84×) *k*-mer coverage peaks supporting the diploid model and indicating sufficient heterozygosity for haplotype-resolved assembly. The Smudgeplot^21^ analysis further supports classifying the Lusitano29 genome as primarily of diploid origin, with *k*-mer pairs dominated by the AB configuration (76% of pairs; AAB 9%, AABB 5%, AAAB 3%) and only a small fraction of higher-copy configurations, indicating limited recent duplication (Fig. 1d). The *k*-mer-based genome size estimate is 9.8% smaller than the flow cytometry estimate, consistent with a widely observed tendency of *k*-mer-based methods to underestimate genome size relative to flow cytometry, particularly in repeat-rich and heterozygous genomes^54,55^.

**Figure 1.**
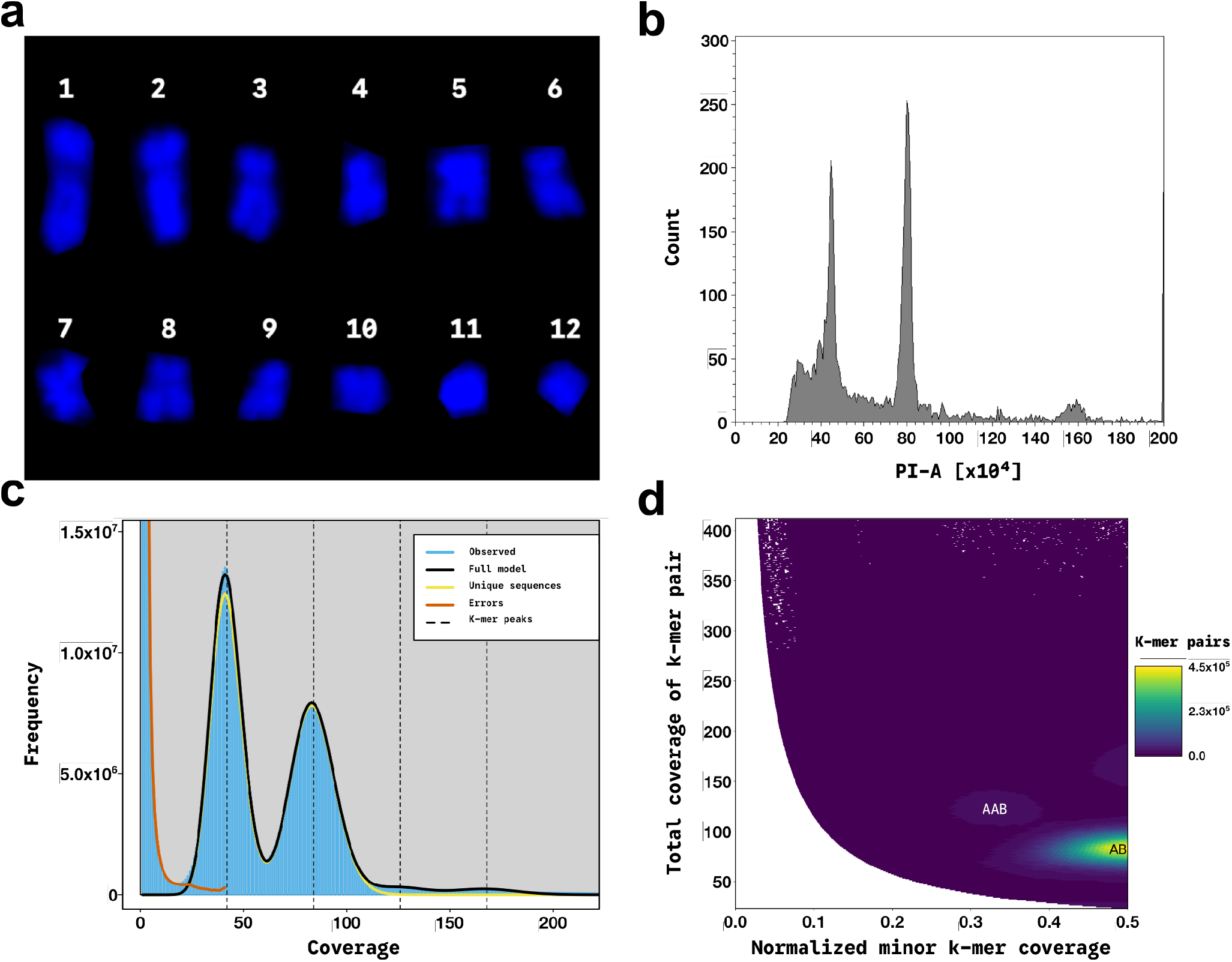
Genome size estimation and k-mer-based genomic profiling of *Lotus pedunculatus* Lusitano29. (a) DAPI-stained karyogram of twelve metaphase chromosomes sorted by size from one independent root-tip preparations of the sequenced genotype. (b) Flow cytometric histogram of propidium iodide fluorescence from nuclei co-chopped with the internal standard *Solanum lycopersicum* (2C = 1.96 pg); left and right peaks correspond to *L. pedunculatus* and the standard, respectively. (c) GenomeScope2 profile of 41-mer frequencies under a diploid model (*p* = 2). The first peak (~ 42 ×coverage) represents k-mers unique to one haplotype (heterozygous), and the second (~ 84×) k-mers shared by both haplotypes (homozygous); the model estimates a haploid genome size of 487.8 Mb, heterozygosity of 1.08%, and a unique-sequence fraction of 66.7%. (d) Smudgeplot of 41-mer pairs; the x-axis shows the normalized minor k-mer coverage B/(A+B) and the y-axis the summed pair coverage. The dominant AB smudge (76% of k-mer pairs) at B/(A+B) *≈*0.5 supports the hypothesis of diploidy for this genotype, with minor contributions from AAB (9%), AABB (5%), and AAAB (3%) configurations.

### CiFi pseudo-paired reads

*In silico* digestion of CiFi concatemers confirmed the expected cleavage frequencies of both restriction enzymes (Table 4; Fig. 2). The 4-bp cutter *Dpn*II produced a broad monomer distribution averaging 17.0 monomers per concatemer, whereas the 6-bp cutter *Hind*III yielded a narrower distribution averaging 3.4 monomers, consistent with more sparse genomic cutting thus resulting in longer monomer sizes^8^. Following the removal of single-monomer concatemers and fragments shorter than 50 bp, combinatorial pairing generated 790.3 million pseudo-paired contacts from *Dpn*II and 10.3 million from *Hind*III, demonstrating the substantial gain in pairwise contact information that distinguishes long-read chromosome conformation capture from short-read Hi-C.

**Table 4.** CiFi concatemer *in silico* digestion and pseudo-pairwise contact generation for the *Dpn*II and *Hind*III libraries of *Lotus pedunculatus* genotype Lusitano29. Concatemers were digested into monomers; monomer counts were recorded after digestion but before filtering. Filtering discarded single-monomer concatemers and monomers shorter than 50 bp.

| Restriction enzyme | Concatemers [M] | Monomers [M] | Read pairs [M] |
| --- | --- | --- | --- |
| *DpnII* | 8.9 | 151.7 | 790.3 |
| *HindIII* | 2.1 | 7.1 | 10.3 |

**Figure 2.**
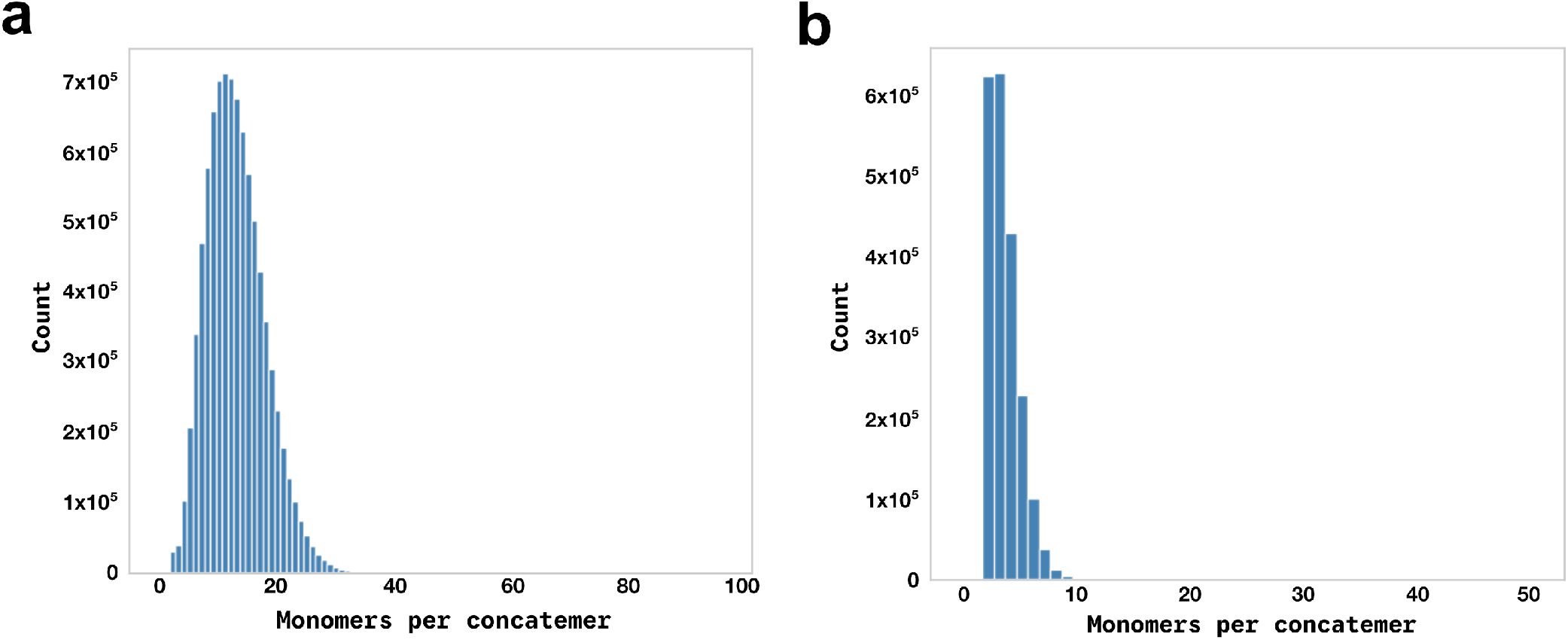
Distribution of monomers per CiFi concatemer from *Lotus pedunculatus* Lusitano29 after *in silico* digestion and filtering (single-monomer concatemers discarded; monomers *<*50 bp removed) for the *Dpn*II (a) and *Hind*III (b) libraries. The frequent cutter *Dpn*II yields a broad distribution peaking at 15–16 monomers per concatemer, whereas the six-cutter *Hind*III yields a narrow distribution peaking at three, reflecting the inverse relationship between restriction site density and monomer length.

### Comparison between manual curation and automated scaffolding

To compare the performance of automated scaffolding against manual curation, the diploid assembly was additionally scaffolded with HapHiC (see Methods). Sequence content and gene-space completeness were identical between automated scaffolding and manual curation (assembly size 991.1 Mb, BUSCO completeness 97.0%, Merqury QV 64.7; Table 5), isolating sequence arrangement into pseudo-chromosomes as the sole difference between approaches. The automated scaffolding was substantially less contiguous at the pseudo-chromosome scale, as HapHiC grouped sequences into only nine clusters against the expected twelve pseudo-chromosomes and failed to place 630 of 668 candidate contigs during reassignment. Consequently, HapHiC produced a scaffold N90 of only 23.6 Mb, compared to 55.4 Mb with manual curation, and a smaller maximum scaffold length (116.1 Mb vs. 133.5 Mb; Fig. 3). The Nx curves further confirm this: the manually curated assembly forms a staircase of twelve chromosome-scale scaffolds terminating in an abrupt drop near *x* = 95, whereas the HapHiC scaffolding falls below the curated curve from *x ≈*50 onward and exhibits a tail of small unplaced sequences beyond *x≈* 80, where *x* denotes the cumulative percentage of total assembly length. A possible explanation for this is that HapHiC was developed for short-read Hi-C rather than CiFi contact data. The curated assembly was therefore retained as the final reference and thus used for all downstream analyses. This comparison is dataset- and configuration-specific and does not constitute a general assessment of HapHiC.

**Table 5.** Assembly metrics for the phased haplotypes 1 and 2 (H1, H2) of *Lotus pedunculatus* Lusitano29 and for the concatenated diploid assembly scaffolded either automatically with HapHiC or by manual curation. Sequence-level statistics (total size, number of sequences, largest sequence, N50, N90) were computed with QUAST, consensus quality and k-mer completeness with meryl/Merqury using the HiFi read set, gene-space completeness with BUSCO in genome mode against the `fabales_odb12.2025-07-01` dataset, and structural and regional quality with CRAQ from read-mapping profiles.

| Metric | Phased contig assembly |  | Diploid scaffolding |  |
| --- | --- | --- | --- | --- |
|  | H1 | H2 | HapHiC | Manual curation |
| **Sequence-level** |  |  |  |  |
| Total assembly size [Mb] | 499.9 | 491.2 | 991.1 | 991.1 |
| Number of sequences | 496 | 172 | 639 | 639<sup>b</sup> |
| Longest sequence [Mb] | 62.3 | 68.2 | 116.1 | 133.5 |
| Sequence N50 [Mb] | 42.7 | 29.9 | 73.6 | 73.8 |
| Sequence N90 [Mb] | 9.8 | 7.5 | 23.6 | 55.4 |
| **Quality & completeness** |  |  |  |  |
| Merqury QV [Phred] | 63.4 | 66.7 | 64.7 | 64.7 |
| Merqury k-mer completeness [%] | 80.8 | 80.8 | 99.4 | 99.4 |
| BUSCO completeness [%]<sup>a</sup> | 96.8 | 96.8 | 97.0 | 97.0 |
| CRAQ structural quality | 98.3 | 99.4 | 100.0 | 100.0 |
| CRAQ regional quality | 99.0 | 98.7 | 99.3 | 99.4 |

<sup>a</sup> Evaluated against the *fabales\_odb12.2025-07-01* database (n=7,843).
<sup>b</sup> During scaffolding, 29 contigs were merged, reducing the 668 input contigs to 639 final curated scaffolds.

**Figure 3.**
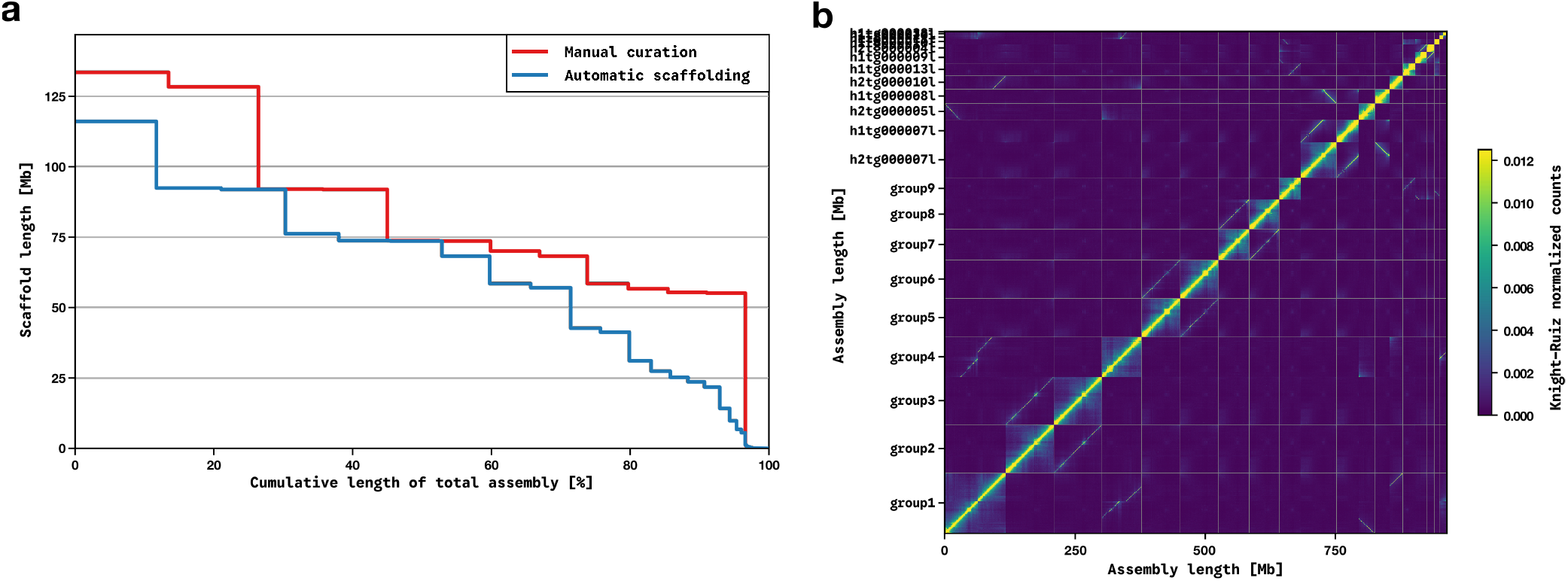
Comparison of automated scaffolding and manual curation of the diploid *Lotus pedunculatus* Lusitano29 assembly. (a) QUAST v.5.3.0 Nx curves for the HapHiC-scaffolded and manually curated assemblies, both derived from the same filtered CiFi alignment file; *x* denotes the cumulative percentage of total assembly length. The manually curated assembly forms a staircase of twelve pseudo-chromosomes terminating in an abrupt drop near *x* = 95, whereas the automated scaffolding shows reduced scaffold lengths from *x≈* 50 and a long tail of small unplaced sequences beyond *x≈* 80 (scaffold N90 of 23.6 Mb vs. 55.4 Mb). (b) Knight–Ruiz normalized CiFi contact map (500 kb bins) of the automated HapHiC scaffolding generated with the `HapHiC plot` module; nine chromosome groups were resolved against the expected twelve, with unplaced contigs visible as individual stripes beyond the main diagonal blocks. The corresponding contact map of the manually curated assembly is shown in Fig. 4a.

**Figure 4.**
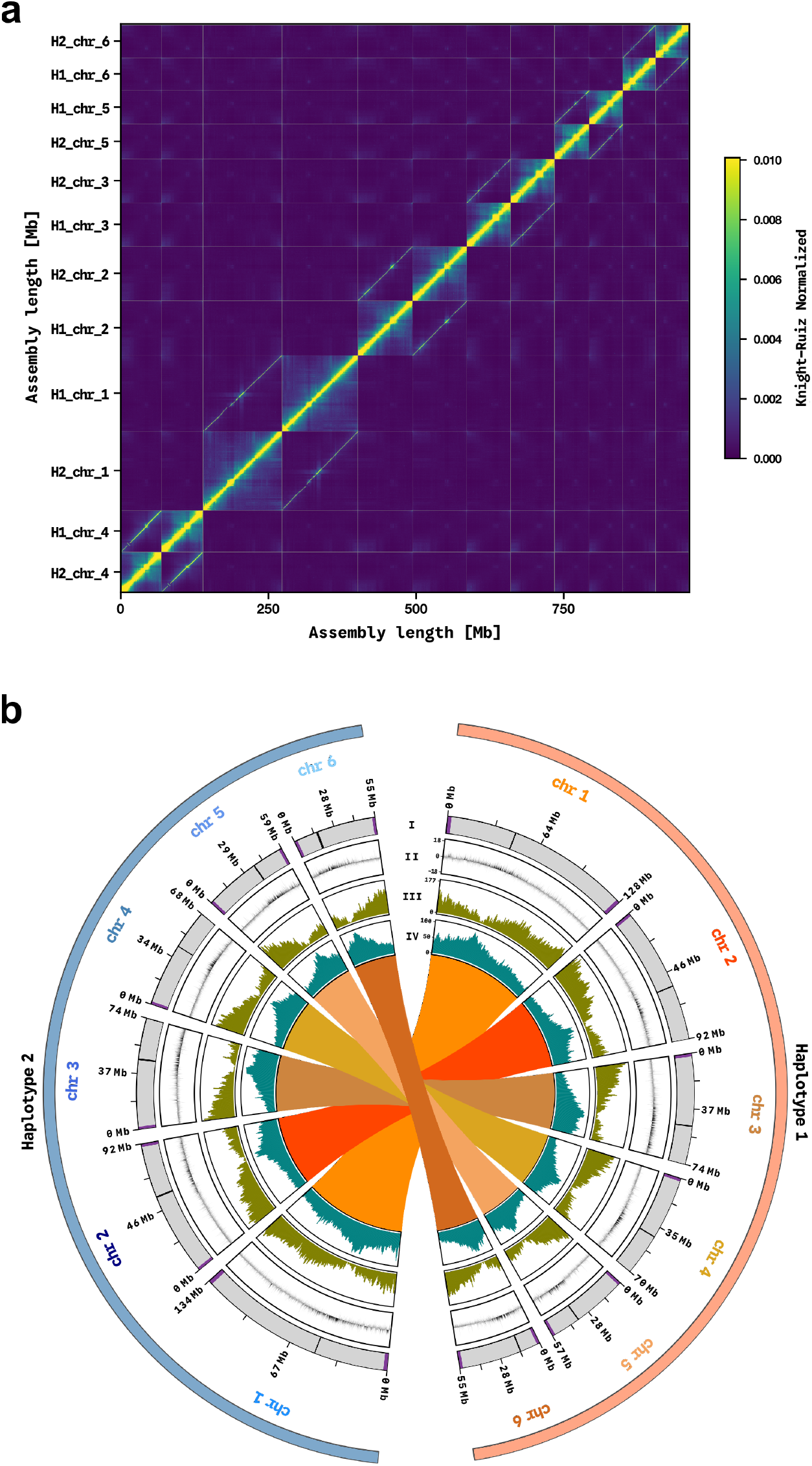
Overview of the haplotype-resolved *Lotus pedunculatus* genotype Lusitano29 genome assembly. (a) Knight–Ruiz normalized CiFi contact matrix of the curated assembly (500 kb bins); bright yellow indicates higher contact frequency. Twelve pseudo-chromosomes appear as distinct squares along the diagonal, ordered so that homologous haplotype 1 (H1) and haplotype 2 (H2) pseudo-chromosome pairs are adjacent; the elevated off-diagonal signal between homologs reflects their spatial proximity in the nucleus. (b) Circos plot of the twelve pseudo-chromosomes, with haplotype 1 (orange) and haplotype 2 (blue) arranged on opposite halves. Tracks from outermost to innermost: I, pseudo-chromosome ideograms with terminal telomeric repeat arrays marked in purple (positions indicative; band sizes are not to scale) and putative centromeric regions in black; II, GC content deviation from the global mean (20 kb windows, 5 kb step; black = above mean, grey = below mean); III, gene density per 1 Mb window (500 kb step); IV, proportion of transposable element bases per 1 Mb window (500 kb step). Inner links connect homologous chromosomes and represent pairwise alignments of at least 2 kb at *≥*80% sequence identity.

### Assembly quality assessment

The curated diploid assembly spans 991.1 Mb with a scaffold N50 of 73.8 Mb and N90 of 55.4 Mb. Per-haplotype contig N50 values were 42.7 Mb (H1) and 29.9 Mb (H2). Of the total assembly, 957.4 Mb (96.6%) was anchored onto the twelve pseudo-chromosomes (476.0 Mb in H1; 481.4 Mb in H2), with the remaining 33.7 Mb distributed across 627 unplaced sequences. Genome-mode BUSCO completeness was 97.0% for the diploid assembly (96.8% per haplotype) and consensus quality was high (Merqury QV 63.4–66.7 per haplotype; 64.7 combined) (Table 5).

Three lines of evidence indicate that the two haplotypes are cleanly phased and structurally balanced. First, Merqury *k*-mer completeness was 80.8% for each individual haplotype but reached 99.4% for the diploid assembly, demonstrating that virtually all read-derived *k*-mers are represented. The lower completeness of individual haplotypes reflects the expected partitioning of heterozygous *k*-mers onto the alternate homolog in a genome with 1.08% heterozygosity, rather than missing sequence. Second, transposable element annotation recovered near-identical repeat fractions in both haplotypes (H1: 53.4% and H2: 54.3%; Table 7), indicating that repeat-rich regions were assembled with equal completeness in both assemblies. Third, CRAQ structural quality scores increased from 98.3% (H1) and 99.4% (H2) when evaluated per haplotype, to 100.0% in the diploid assembly. Because misjoins lack read support across both haplotypes and persistently generate clipped alignments^35^, the loss of structural penalties upon combining both haplotypes confirms that read clipping was driven by heterozygous structural variants – such as indel polymorphisms segregating between the homologs – rather than assembly errors. Furthermore, regional quality scores remained exceptionally high across all evaluations (99.0% for H1, 98.7% for H2, and 99.4% diploid), consistent with the capacity of long HiFi reads to maintain high mapping fidelity across complex repetitive regions^35^.

### Validation of gene and transposable element annotations

The final annotation pipeline yielded 38,069 genes (51,148 mRNAs) in H1 and 36,484 genes (49,587 mRNAs) in H2 (Table 6). When evaluated using the longest isoform per gene via AGAT v0.8.0^41^ to avoid isoform-inflated duplication, protein-mode BUSCO completeness of the initial EviAnn annotations reached 96.3% for both haplotypes (duplication rates of 6.2% in H1 and 5.8% in H2). Following complementation with non-overlapping Helixer^40^ *ab initio* gene models, completeness increased to 96.5% across both haplotypes with only minor increases in duplication (6.7% and 6.2%, respectively), confirming that the incorporated models captured previously unannotated gene space rather than redundant features. In contrast to the annotated gene content, the repetitive sequences in Lusitano29 are predominantly composed of LTR retrotransposons (Table 7), with Gypsy (H1: 13.5%; H2: 14.5%) and Copia (H1: 11.3%; H2: 11.5%) elements representing the most abundant superfamilies in both haplotypes.

**Table 6.** Structural annotation summary statistics for the nuclear gene sets of haplotypes 1 and 2 (H1, H2) of *Lotus pedunculatus* Lusitano29 at three stages of the annotation pipeline. EviAnn: evidence-based structural annotation integrating Iso-Seq transcripts, *Lotus japonicus* coding sequences, and Hologalegina RefSeq proteins. Helixer: *ab initio* predictions from the deep-learning model, used only as a complementation source. Final: the published annotation, generated by adding Helixer gene models only where they did not overlap any existing EviAnn gene; EviAnn models were otherwise retained unchanged, so the difference between the EviAnn and Final gene counts (10,255 genes added to H1; 8,996 to H2) represents the non-overlapping Helixer models incorporated. Mean values are computed across all annotated features of each class.

| Metric | EviAnn |  | Helixer |  | Final |  |
| --- | --- | --- | --- | --- | --- | --- |
|  | H1 | H2 | H1 | H2 | H1 | H2 |
| Total number of genes | 27,814 | 27,488 | 35,360 | 33,881 | 38,069 | 36,484 |
| Total number of mRNAs | 40,893 | 40,591 | 35,360 | 33,881 | 51,148 | 49,587 |
| Mean mRNAs per gene | 1.5 | 1.5 | 1.0 | 1.0 | 1.3 | 1.4 |
| Mean gene length [bp] | 4,329 | 4,391 | 3,748 | 3,781 | 3,778 | 3,839 |
| Mean mRNA length [bp] | 4,617 | 4,670 | 3,748 | 3,781 | 4,149 | 4,213 |
| Mean CDS length [bp] | 1,400 | 1,401 | 1,218 | 1,221 | 1,277 | 1,280 |
| Mean exons per mRNA | 6.0 | 6.1 | 5.5 | 5.6 | 5.6 | 5.7 |
| Mean exon length [bp] | 318 | 316 | 304 | 302 | 311 | 309 |

**Table 7.**
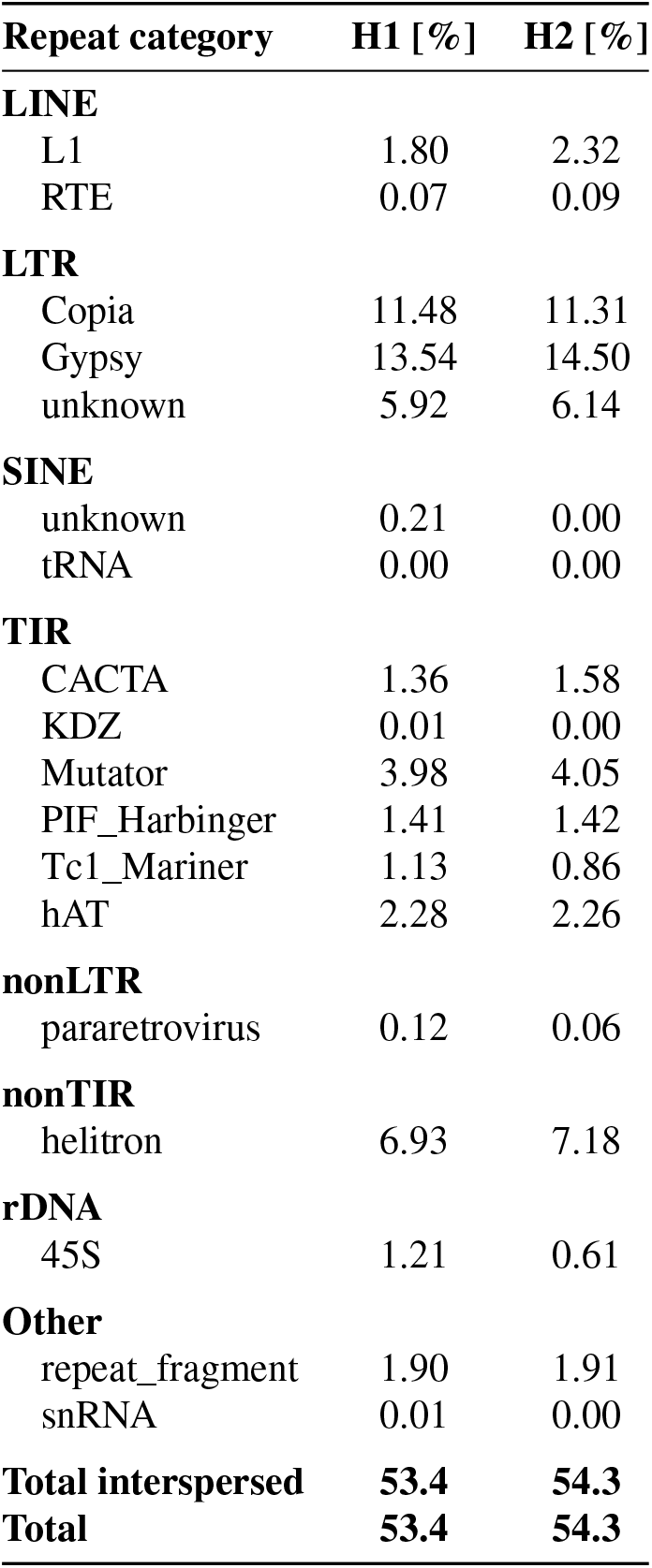
Transposable element composition (repeat categories) of *Lotus pedunculatus* Lusitano29 haplotypes 1 and 2 (H1, H2), annotated with EDTA using a *de novo* repeat library built by the default pipeline. Values give the proportion of each assembly (in percent) assigned to each repeat category. ‘Total interspersed’ excludes tandem and low-complexity repeats; ‘Total’ includes all annotated repeat content.

| Repeat category | H1 [%] | H2 [%] |
| --- | --- | --- |
| **LINE** |  |  |
| L1 | 1.80 | 2.32 |
| RTE | 0.07 | 0.09 |
| **LTR** |  |  |
| Copia | 11.48 | 11.31 |
| Gypsy | 13.54 | 14.50 |
| unknown | 5.92 | 6.14 |
| **SINE** |  |  |
| unknown | 0.21 | 0.00 |
| tRNA | 0.00 | 0.00 |
| **TIR** |  |  |
| CACTA | 1.36 | 1.58 |
| KDZ | 0.01 | 0.00 |
| Mutator | 3.98 | 4.05 |
| PIF_Harbinger | 1.41 | 1.42 |
| Tc1_Mariner | 1.13 | 0.86 |
| hAT | 2.28 | 2.26 |
| **nonLTR** |  |  |
| pararetrovirus | 0.12 | 0.06 |
| **nonTIR** |  |  |
| helitron | 6.93 | 7.18 |
| **rDNA** |  |  |
| 45S | 1.21 | 0.61 |
| **Other** |  |  |
| repeat_fragment | 1.90 | 1.91 |
| snRNA | 0.01 | 0.00 |
| **Total interspersed** | **53.4** | **54.3** |
| **Total** | **53.4** | **54.3** |

### Structural validation

Telomeric repeats were identified at 19 of 24 pseudo-chromosome termini. All twelve pseudo-chromosomes carry telomeric repeats at least one terminus and seven carry repeats at both ends (Fig. 4b). Putative centromeric regions were inferred for all twelve pseudo-chromosomes from the co-localization of interstitial telomeric repeat enrichment, elevated transposable element density and reduced gene density (Fig. 4b). These regions are considered putative because interstitial telomeric repeats have not been experimentally demonstrated to mark centromeres in *Lotus*. All-against-all synteny analysis between the H1 and H2 assemblies revealed strictly conserved one-to-one collinearity across all six chromosome pairs, with no evidence of large-scale non-homologous translocations or chimeric scaffolding errors. Pairwise collinear blocks between homologous pseudo-chromosomes demonstrate near-complete structural concordance between the two phased haplotypes (Fig. 4b). The final CiFi contact map of the curated assembly shows twelve pseudo-chromosomes with contact signal concentrated along the diagonal and the expected elevated inter-homolog signal between adjacent homologous pseudo-chromosomes (Fig. 4a).

## Code Availability

Scripts and commands used for quality control, assembly, scaffolding, annotation and technical validation are available at https://github.com/a-pettersson/MPB-GA-drLotPedu.

## Author Contributions

A.P.: Conceptualization, Methodology, Software, Validation, Formal Analysis, Investigation, Data Curation, Writing – Original Draft, Writing – Review & Editing, Visualization. Y.C.: Methodology, Writing – Review & Editing, Supervision. T.D.: Methodology, Writing – Original Draft, Writing – Review & Editing. P.N.: Conceptualization, Methodology, Writing – Review & Editing. D.K.: Methodology, Investigation, Writing – Original Draft, Writing – Review & Editing, Visualization. B.S.: Conceptualization, Resources, Writing – Review & Editing, Supervision, Project Administration, Funding Acquisition. R.K.: Conceptualization, Resources, Writing – Review & Editing, Supervision, Project Administration, Funding Acquisition. All authors read and approved the final manuscript.

## Funding

This study was supported by the BELIS project (“Breeding European Legumes for Increased Sustainability”), which has received funding from the European Union’s Horizon Europe research and innovation programme under grant agreement No. 101081878. D.K. was supported by the ERDF Programme Johannes Amos Comenius (project TowArds Next GENeration Crops, reg. no. CZ.02.01.01/00/22_008/0004581).

This study was funded by the European Union. Views and opinions expressed are however those of the author(s) only and do not necessarily reflect those of the European Union or the European Research Executive Agency. Neither the European Union nor the granting authority can be held responsible for them.

## Competing interests

The authors declare no competing interests.

## Acknowledgements

The authors thank the Genetic Diversity Center and Ingrid Stoffel-Studer at ETH Zurich for their help in producing the sequencing data. We also thank Verena Knorst for greenhouse and plant logistical assistance and Selina Steerup Moore and Carsten Stefan Malisch from Aarhus University for providing the Lusitano29 genotype.

## Notes

https://www.ebi.ac.uk/ena/browser/view/PRJEB120729

### Competing Interest Statement

The authors have declared no competing interest.

